# A computational panel of pathological RAS mutants with implications for personalized medicine and genetic medicine

**DOI:** 10.1101/153726

**Authors:** Chelsey Kenney, Edward C. Stites

**Affiliations:** Clinical Translational Research Division, Translational Genomics Research Institute, Phoenix, Arizona; School of Biological and Health Systems Engineering, Arizona State University, Tempe, Arizona; Integrative Biology Laboratory, Salk Institute for Biological Studies, La Jolla, California

## Abstract

The RAS proteins (KRAS, NRAS, and HRAS) play important roles in multiple diseases. This includes many types of cancer and the developmental syndromes collectively referred to as the RASopathies. There are many different RAS mutants that are found to drive these diseases. Mutant-to-mutant differences pose a challenge for personalized medicine. To investigate this problem, we extend our previously developed model of oncogenic RAS mutants to a total of 16 oncogenic mutants. We also extend our model to RASopathy associated mutants using data for 14 such RAS mutants. The model finds that the known biochemical defects of these mutants are typically sufficient to explain their elevated levels of RAS signaling. In general, our analysis finds that the oncogenic mutants are stronger than the RASopathy mutants. However, the model suggests that RAS signal intensities are spanned by the pathological variants; there does not appear to be a perfect separation between cancer promoting and developmental syndrome promoting mutants. Analysis of the panel also finds that the relative strengths of pathological RAS mutants is not absolute, but rather can vary depending on context. We discuss implications of this finding for personalized cancer medicine and for medical genetics. As genomics permeates clinical medicine, computational models that can resolve mutant specific differences, like the one presented here, may be useful for augmenting clinical thinking with their ability to logically translate biochemical knowledge into system level outputs of perceived clinical relevance.

## INTRODUCTION

Multiple human diseases are associated with *RAS* gene mutations that encode mutated RAS proteins. Somatic mutations to the *RAS* genes are common in many cancers, including lung, colon, and pancreatic cancer [1-3]. Germline mutations to the *RAS* genes are found in several developmental syndromes, including Noonan syndrome [4] and Costello syndrome [5]. These syndromes are members of a group of developmental syndromes with overlapping phenotypes that are collectively referred to as RASopathies because the causative mutations occur within RAS signaling pathway genes [6].

Many different point mutations to the *RAS* genes have been found within patients with these diseases. The presence of multiple mutations of similar, but non-identical, function associated with a disease poses multiple problems for genomic medicine. For example, different mutations to the same gene may respond differently to the same drug [7]. Genetic and genomic medicine will progress slowly if each mutation must be considered separately. For example, it will be more difficult to acquire large cohorts of patients to study individual mutations compared to studying patients with functionally equivalent mutations. Methods to better anticipate mutant behaviors and to better extrapolate between mutants are needed. Clinicians and scientists have few tools to investigate this problem. Additionally, there are limited examples on which to build intuition. This lack of knowledge and intuition poses a significant challenge to the advancement of personalized medicine.

We have previously developed a mathematical model of the processes that regulate RAS nucleotide binding [8, 9]. Simulations of this model have proven effective at estimating the proportion of mutant RAS in the “on” state responsible for pathological signaling. This has also allowed for many different questions about mutant-driven RAS signaling to be investigated *in silico* and then experimentally [8, 10].

With the demonstrated challenges of intuiting how pathological mutations behave, and the demonstrated success of the RAS model, we decided to apply our model to the problem of studying multiple RAS mutations. Our goal was to develop intuition that could apply to problems in personalized medicine and genetic medicine. We extend our RAS model to a total of 16 oncogenic RAS mutants and to 14 RASopathy RAS mutants. We find patterns of activation of the model to be consistent with known patterns of activation, such as oncogenic mutants tending to be stronger than RASopathy mutants, and the known strong (and weak) mutants in each class being among the strongest (and weakest). This demonstrates that known biochemical properties of these mutants, when analyzed with our model, tend to be sufficient to explain the pathological increase in signal. Interestingly, we find the relative strengths of RAS mutants is not fixed, but can rather vary between conditions. In other words, if mutant A is stronger than mutant B in condition X, it cannot be assumed that mutant A is stronger than mutant B in condition Y. We discuss implications of this in cancer and in the RASopathies.

## MODEL

The model is based on the well-known mechanisms of RAS signaling. RAS is commonly considered to be a switch. RAS bound to GDP is considered to be in the “off” state, and RAS bound to GTP is considered to be in the “on” state. Whether RAS is in the “on” state and bound to GTP, or in the “off” state and bound to GDP is controlled by several different processes. Individual RAS proteins cycle between “on” and “off” states. RAS mutations that promote disease also cycle between “on” and “off” states, but disease promoting mutations tend to spend a larger proportion of time in the “on” state compared to a cell with all wild-type RAS. The disease promoting mutations typically alter the biochemical rate constants for one or more of the processes that regulate RAS signaling to result in a shifting of the dynamic equilibrium toward the “on” state.

The details of the model have been described extensively in our previous publications [8-13]. Briefly, the model includes RAS GTPases, GAPs (negative regulators of RAS), GEFs (positive regulators of RAS) and Effector proteins with which RAS directly interacts. GAPs, GEFs, and Effectors are all grouped into one pool each; there are two pools of RAS in the model (a wild-type pool and a mutant pool) (Figure 1). The entire model is specified with nine coupled, nonlinear, ordinary differential equations. Each equation corresponds to the rate of change of one possible state of RAS (RASGDP, RASGTP, RAS nucleotide free, RASGTP-Effector) for both wild-type and mutant RAS, as well as a pool of free Effector proteins (Figure 2).

**Figure 1:**
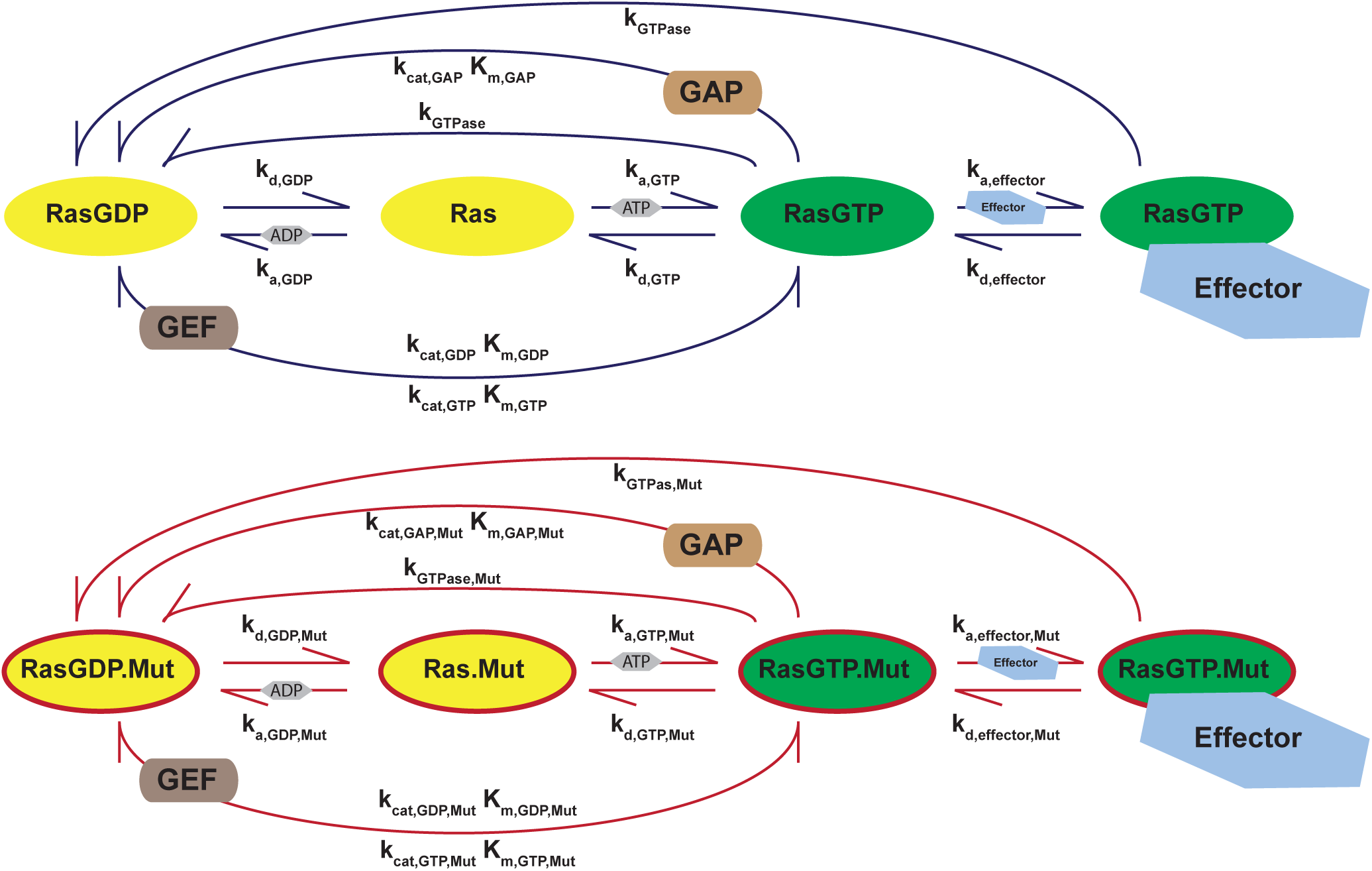
Schematic of RAS signal regulation for wild-type and mutant RAS. The four states of RAS proteins in the model (bound to GDP, bound to GTP, bound to no nucleotide, or bound to GTP and Effector) and their paths between states with biochemical parameters and partners indicated.

**Figure 2:**
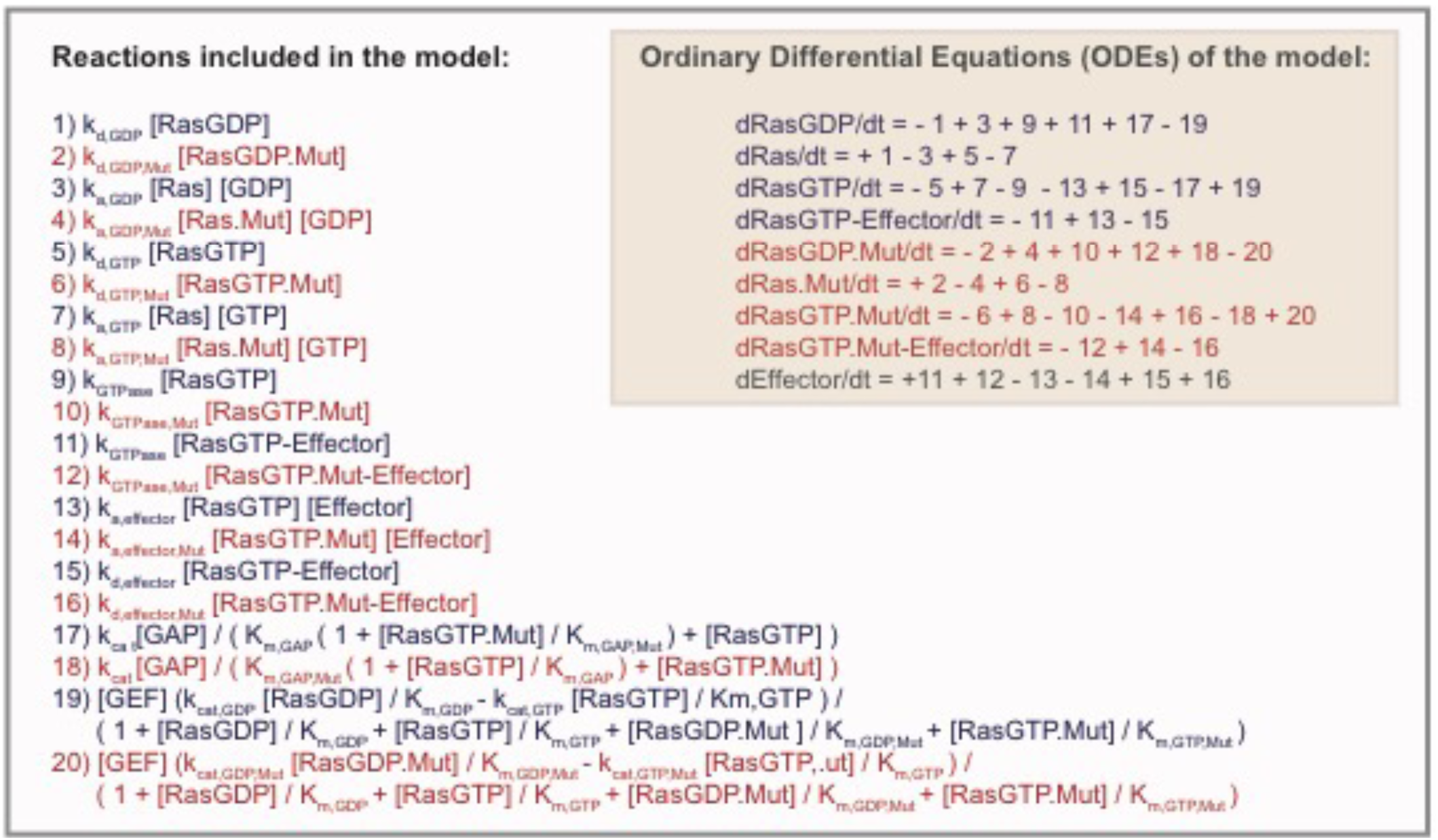
Equations of the RAS model. The reactions indicated in Figure 1 are individually specified by mass-action kinetics or Michaelis-Menten kinetics (for GEF and GAP reactions). The ordinary differential equation model is built from the individual reaction specifications.

Within our model, a RAS mutant is modeled by the choice of rate constants used to characterize the reactions involved in the regulation of RAS signaling. We obtain these parameter values from the experimental literature (Table I). As the value of wild-type rate constants can vary between different studies that measure the same reaction, we focus upon the ratio of change between mutant and wild-type rate constants within a single study. This ratio characterizes the change in value of the mutant for that parameter, and is multiplied by our previously set value for the corresponding wild-type parameter. In cases where we could not find that a parameter has been measured, we assume no change from wild-type RAS. As such, the values in this panel can be considered estimates based on currently available data. The analysis of these computational mutants allows us to both assess whether known biochemical changes are consistent with observed changes in total RASGTP (a system property). We can also use this panel as a study set to investigate consequences of variation between mutants.

**Table I.** Parameters used to model the expanded pool of computation pool of pathological RAS mutants. Table presents factors by which corresponding wild-type parameter changes for the mutant of interest. Factor is determined by comparing ratios of mutant to wild-type parameter in the indicated reference; truncated decimal representation presented in table. When the property was not measured, wild-type (WT) parameter values are utilized. NC: no change. For incompletely determined parameters, such as when K_d_ had measured (rather than k_a,Eff_ and k_d,Eff_), we consider k_a,Eff_, K_M,GDP,GEF_, and K_M,GTP,GEF_ to be unchanged and apply the change to the equilibrium constant to K_d,Eff_ or K_cat,GDP,GEF._ Because of thermodynamic equivalence between intrinsic and GEF mediated exchange, all parameters in these processes are not independent. The parameter k_cat,GTP,GEF_ was designated function of the intrinsic and GEF mediated exchange parameters to ensure thermodynamic validity. Mutants for the model first described here are specified in bold.

| | $k_{\text{GTPase}}$ | | $k_{d, \text{GDP}}$ | | $k_{d, \text{GTP}}$ | | $k_{a, \text{GDP}}$ | | $k_{a, \text{GTP}}$ | | $k_{\text{cat, GAP}}$ | | $K_{M, \text{GAP}}$ | | $k_{d, \text{Eff}}$ | | $k_{\text{cat, GDP, GEF}}$ | |
| --- | --- | --- | --- | --- | --- | --- | --- | --- | --- | --- | --- | --- | --- | --- | --- | --- | --- | --- |
| G12V | 0.15 | [37] | 0.31 | [37] | 0.80 | [37] | 2.27 | [37] | 4.14 | [37] | NC |  | WT |  | 0.44 | [38] | WT |  |
| G12D | 0.40 | [37] | 0.48 | [37] | 5.00 | [37] | 1.37 | [37] | 3.43 | [37] | NC |  | WT |  | WT |  | WT |  |
| **G12P** | 1.54 | [21] | 0.70 | [21] | 1.18 | [21] | WT |  | WT |  | 0.01 | [39] | WT |  | WT |  | WT |  |
| **G12R** | 0.14 | [40] | 0.47 | [40] | 0.05 | [40] | WT |  | WT |  | NC |  | WT |  | WT |  | WT |  |
| G12C | 0.72 | [41] | WT |  | WT |  | WT |  | WT |  | NC |  | WT |  | 1.20 | [41] | WT |  |
| **G13V** | 0.46 | [42] | 79 | [42] | 6.00 | [42] | WT |  | WT |  | NC |  | WT |  | WT |  | WT |  |
| **G13S** | 1.14 | [42] | 2.91 | [42] | 1.00 | [42] | WT |  | WT |  | NC |  | WT |  | WT |  | WT |  |
| G13D | 0.14 | [41] | 3.00 | [43] | 3.00 | [43] | WT |  | WT |  | NC |  | 100 | [44] | WT |  | WT |  |
| Q61L | 0.02 | [19] | 2.46 | [19] | 1.07 | [19] | WT |  | WT |  | NC |  | 0.06 | [19] | 0.78 |  | WT |  |
| Q61K | 0.02 | [19] | 1.40 | [19] | 0.60 | [19] | WT |  | WT |  | NC |  | 57.14 | [19] | WT |  | WT |  |
| Q61W | 0.07 | [19] | 1.00 | [19] | 0.80 | [19] | WT |  | WT |  | NC |  | 2.86 | [19] | WT |  | WT |  |
| Q61R | 0.02 | [19] | 1.14 | [19] | 0.45 | [19] | WT |  | WT |  | NC |  | 42.86 | [19] | WT |  | WT |  |
| Q61H | 0.09 | [19] | 1.48 | [19] | 0.61 | [19] | WT |  | WT |  | NC |  | 0.71 | [19] | WT |  | WT |  |
| Q61P | 0.08 | [19] | 1.05 | [19] | 1.27 | [19] | WT |  | WT |  | WT |  | 1.43 | [19] | WT |  | WT |  |
| **A59T** | 0.24 | [40] | 0.56 | [40] | 1.22 | [40] | WT |  | WT |  | NC |  | WT |  | WT |  | WT |  |
| **10G11** | 8.46 | [45] | 5.46 | [45] | 2.36 | [45] | WT |  | WT |  | NC |  | WT |  | WT |  | WT |  |
| **E153V** | 0.47 | [24] | 1.67 | [24] | 1.67 | [24] | 0.08 | [24] | 0.10 | [24] | 0.87 | [24] | 2.68 | [24] | 2.68 | [24] | 2.22 | [24] |
| **F156L** | 0.85 | [24] | 63 | [24] | 63 | [24] | 0.16 | [24] | 0.19 | [24] | 0.03 | [24] | 22 | [24] | 22 | [24] | 33 | [24] |
| **G60R** | 0.08 | [24] | 1.33 | [24] | 1.33 | [24] | 0.19 | [24] | 0.80 | [24] | NC | [24] | 51 | [24] | 51 | [24] | NC | [24] |
| **K5N** | 1.01 | [24] | 1.33 | [24] | 1.33 | [24] | 0.06 | [24] | 0.06 | [24] | WT | [24] | 0.73 | [24] | 0.73 | [24] | 1.22 | [24] |
| **P34L** | 0.99 | [24] | 2.67 | [24] | 2.67 | [24] | 0.38 | [24] | 0.54 | [24] | NC | [24] | 125 | [24] | 125 | [24] | 1.56 | [24] |
| **P34R** | 0.96 | [24] | 2.00 | [24] | 2.00 | [24] | 0.56 | [24] | 0.69 | [24] | NC | [24] | 108 | [24] | 108 | [24] | 0.67 | [24] |
| **Q22E** | 1.11 | [24] | 26 | [24] | 26 | [24] | 0.08 | [24] | 0.11 | [24] | 0.03 | [24] | 6.41 | [24] | 6.41 | [24] | 2.22 | [24] |
| **Q22R** | 0.69 | [24] | 1.33 | [24] | 1.33 | [24] | 0.15 | [24] | 0.16 | [24] | 0.04 | [24] | 2.32 | [24] | 2.32 | [24] | 2.22 | [24] |
| **T58I** | 0.58 | [24] | 4.33 | [24] | 4.33 | [24] | 0.81 | [24] | 1.08 | [24] | 0.88 | [24] | 6.32 | [24] | 6.32 | [24] | 1.67 | [24] |
| **V14I** | 1.08 | [24] | 29 | [24] | 29 | [24] | 0.11 | [24] | 0.22 | [24] | WT | [24] | 6.18 | [24] | 6.18 | [24] | 19 | [24] |
| **DupGlu37** | 0.27 | [25] | 2.35 | [25] | 8.47 | [25] | 1.06 | [25] | 1.06 | [25] | WT |  | 266 | [25] | 154 | [25] | 0.38 | [25] |
| **Y71H** | 2.28 | [28] | 1.44 | [28] | 1.44 | [28] | WT |  | WT |  | 2.08 | [28] | WT |  | 0.26 | [28] | 0.50 | [28] |
| **K147E** | 0.78 | [28] | 150 | [28] | 150 | [28] | WT |  | WT |  | 0.75 | [28] | WT |  | 2.17 | [28] | 0.42 | [28] |
| **K117R** | WT | [46] | 134 | [46] | 134 | [46] | WT |  | WT |  | WT | [46] | WT |  | WT |  | WT |  |

We assume that all hot-spot mutations to oncogenic RAS proteins have no appreciable change in GTP hydrolysis upon binding to GAPs [14]. If only one nucleotide dissociation rate constant was measured in an experimental study, we assume an equivalent change in dissociation for the other nucleotide. If an enzymatic deficiency was measured for a RASopathy mutant and it was not specifically attributed to k_cat_ or K_m_, we apply the change to k_cat._ For Noonan syndrome mutants, which generally had significantly decreased capacity to bind effectors, we assumed a similar deficiency in binding to GAPs by increasing the GAP-RAS K_m_ by same factor that the Effector-RAS K_d_ increased.

## RESULTS

### Expansion of the model to additional oncogenic mutants

Our original RAS model focused on the oncogenic G12D and G12V mutants, as well as fast-cycling F28L mutant [8]. To study problems pertaining to specific RAS mutants important to cancer, we have since extended our model to include G13D, Q61L, Q61K, Q61W, Q61H, Q61P, and Q61R [15] and G12C. We here extend our model to an additional 6 oncogenic mutations: G12R, G12P, G13S, G13V, A59T, and 10dupG, yielding a total of 16 different oncogenic mutants

We considered the case where all of the RAS in the cell is mutated (Figure 3A,B). Simulations of the mathematical model were used to find the amount of RASGTP and RASGTP-Effector complex (the active signaling complex) that would occur for each of these mutations when all of the RAS in the modeled network is mutated. We use this case where all of the RAS is mutant to assess the “intrinsic strength” of the mutant. Simulations find that the available data, when analyzed with our model, are sufficient to result in elevated levels of RASGTP and RASGTP-Effector signaling complex for all of the oncogenic mutants. The simulations also demonstrate that known RAS biochemistry suggests different oncogenic mutants have different strengths, i.e. result in defferent levels of active RASGTP and productive RASGTP-Effector signaling complexes. In recent years, there have been many studies that demonstrate different oncogenic mutants have different behaviors and/or different associations with cancer [16-18]. We hypothesize that differences in the strength of the different mutants, as suggested by our modeling, may underlie some of these different biological behaviors.

**Figure 3:**
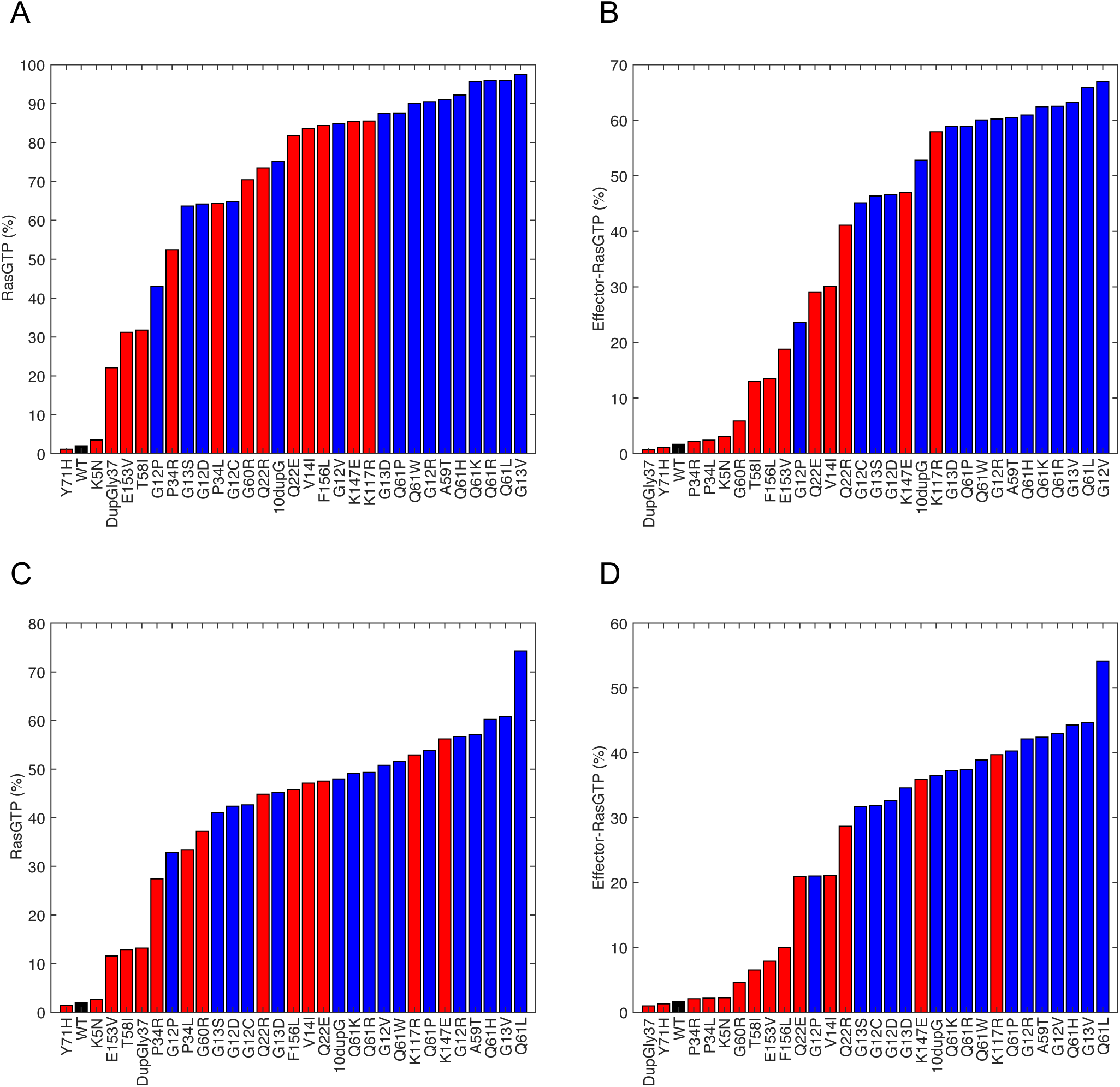
Levels of RAS signal for the computational panel of pathological RAS mutants. Simulations of the model were used to find level of RAS signal for 30 different pathological RAS mutants. The intrinsic strength of each mutant is presented in A) for RASGTP signal and B) RASGTP-Effector complex by simulating 100% of total RAS mutated. The strengths of each mutant in conditions where one allele is mutated and one allele is wild-type is in C) for RASGTP and D) for RASGTP-Effector. RASopathy mutants: red; oncogenic mutants: blue.

In general, a RAS mutation occurs at one of the two RAS alleles. We therefore also considered the case where 50% of the RAS is mutated, and 50% of the RAS is wild-type (Figure 3C,D). In general, the trends are similar, although the magnitude of activation is less. In both cases, we note that known strong mutations and known weak mutations are at the extremes of the oncogenic mutations. The Q61L mutation, for example, is generally considered to be a highly activating RAS mutation, which is consistent with its observed transformation activity in multiple comparative studies [19, 20]. Additionally, G12P mutation has long been considered an exceptionally weak hot-spot mutation [21]. Of note, the G12P mutation is only noted one time in the current release of the COSMIC database for NRAS, and not once for KRAS and HRAS [22]. According to model-based analysis of existing biochemical data characterizing, the Q61L mutant is found among the strongest of the mutants studied and the G12P mutant is found among the weakest of the hot-spot mutants studied.

### Expansion of the panel to RASopathy RAS mutants

We also incorporated parameters for 14 germline RAS mutations observed in Noonan Syndrome (Table I). These mutations that have been observed in patients with a RASopathy include: E153V, F156L, G60R, K5N, P34L, P34R, Q22E, T58I, V14I, DupGly37, Y71H, K147E, K117R, and Q22R. We used simulations of the mutants at different levels of relative expression to evaluate these mutants, as we did for the oncogenic mutants. The intrinsic strength of these mutants (100% mutant) is presented in Figure 3A,B. It is generally believed that RASopathy mutants like these tend to be intrinsically less strong than the oncogenic mutants, and this has been demonstrated experimentally [23-25]. Our model finds that RASopathy mutants tend to be weaker than oncogenic mutants, although this was not universally true. We also considered 50% mutant, 50% wild-type for the reasons discussed above (Figure 3C,D). The same trend with RASopathy mutants tending to be weaker than oncogenic mutants was also observed.

### Modeling suggests RASopathy mutants have less correlation between levels of RASGTP and productive signaling complexes than oncogenic mutants

A comparison of RASopathy mutants and oncogenic mutants is notable for several reasons. First, the RASopathy mutants clearly tend to be weaker than oncogenic mutants (Figure 3). This is consistent with known experimental data comparing these mutants [4, 24] and is consistent with expert opinion on these mutants [6, 23]. That the model naturally reproduces this pattern via incorporation of measured biochemical rate constants that were published after our model was developed and published highlights the quality of information in the RAS mutation field and its logical consistency.

It is worth noting that levels of RASGTP are not as clearly segregated between oncogenic and RASopathy mutants as they are for RASGTP-Effector (Figure 3B,D). The explanation between this discrepancy is that many RASopathy mutants have been described to have impaired binding to effector proteins. That is, many RASopathy mutants are predicted to be highly bound to GTP, but their ability to signal is impaired due to the reported decrease in the ability to bind to effectors [24]. This can also be seen when levels of RASGTP and RASGTP-Effector are compared for the different mutants in these different conditions (Figure 4). Visualizing the relationship in this manner demonstrates that the RASopathy associated mutants tend to fall off of the diagonal much more than the oncogenic mutants. Of note, there is generally a good correlation between levels of RASGTP between high and low levels of mutation, and between levels of RASGTP-Effector complex between high and low levels of mutation (Figure 5).

**Figure 4:**
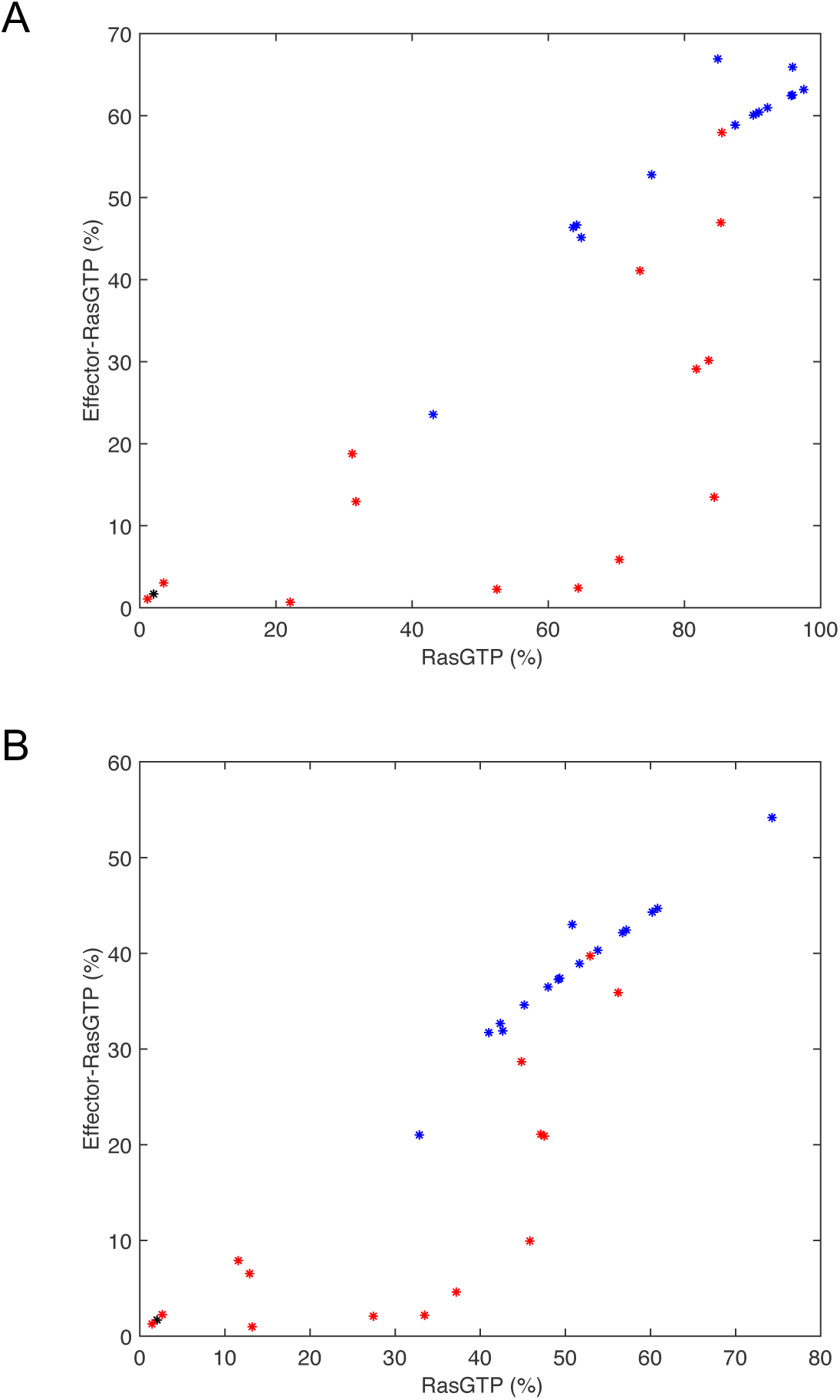
Relationship between RAS loading with GTP and RAS binding to Effector. A) Presents each of the 30 mutants’ values of RASGTP and RASGTP-Effector for intrinsic strength of each mutant (data from Figure 3A,B). B) Presents each of the 30 mutants’ values of RASGTP and RASGTP-Effector for conditions of 50% mutated and 50% wild-type. RASopathy mutants: red; oncogenic mutants: blue.

**Figure 5:**
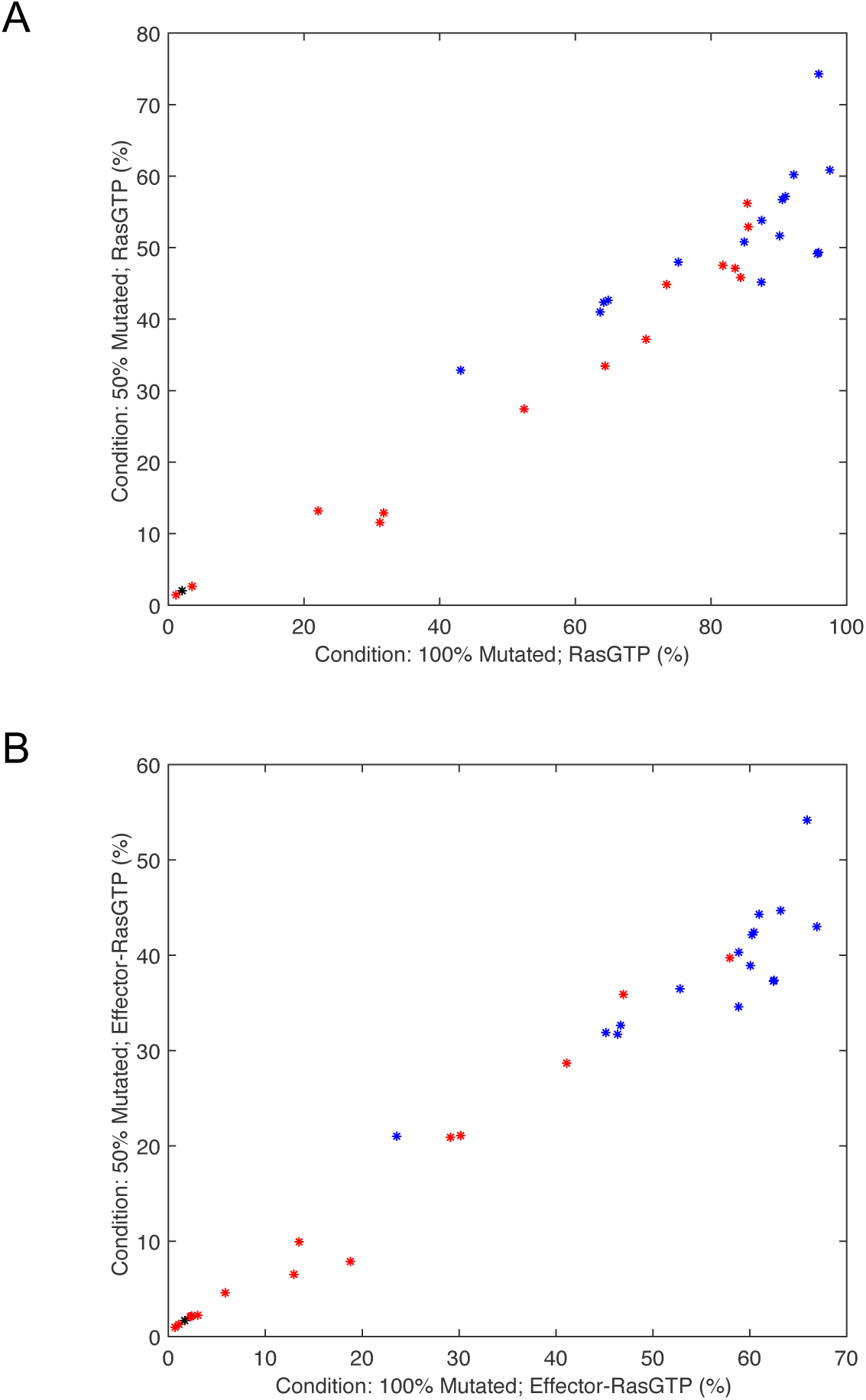
Relationship between RAS signal at different proportions of total RAS mutated. A) Comparison of RASGTP levels for conditions of 100% mutated and 50% mutated. B) Comparisons of RASGTP-Effector levels for conditions of 100% mutated and 50% mutated. RASopathy mutants: red; oncogenic mutants: blue.

### Modeling finds a spectrum of strengths for pathological RAS mutants

Within a cell, KRAS, NRAS, and HRAS proteins may all be expressed [26, 27]. Even when a mutation is present, there is still wild-type RAS within the cell. For example, when KRAS is mutated, NRAS and HRAS are generally present and in their wild-type (non-mutated) form. Therefore, in a cell with a RAS mutation, only a fraction of total RAS is mutated. We now consider total RAS signal for different proportions of total RAS mutated and 50% RAS mutated. Figure 6 shows absolute signal intensities for all of the RAS mutants in the panel, color coded to distinguish oncogenic mutants from RASopathy mutants. There is a trend that mutations identified as “RASopathy” mutants tend to be less activating than mutations identified as “oncogenic”. However, there is a clear overlap and a lack of a clear separation. This suggests that there is a spectrum of signal strengths and not a clear segregation into “strongly activating” and “moderately activating” mutations, as it might appear from experimental studies that are typically limited to a small number of mutants in a limited number of conditions.

**Figure 6:**
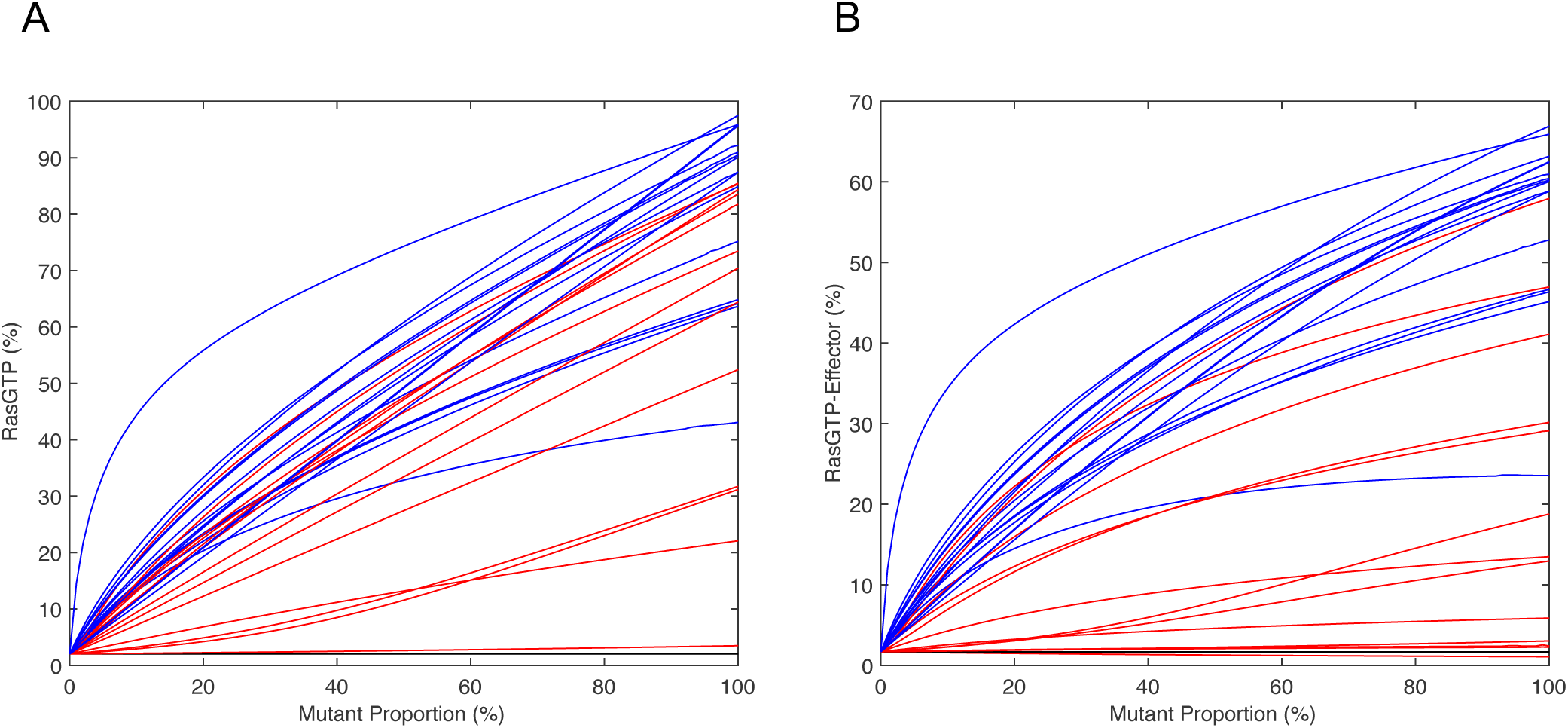
RAS signal as a function of the proportion of total RAS mutated for the computational panel of pathological RAS mutants. A) RASGTP levels. B) RASGTPEffector levels. RASopathy mutants: red; oncogenic mutants: blue.

### The relative strengths of RAS mutants are not absolute

We further investigated the strengths of RAS mutants in more detail. First, we focus on the oncogenic RAS mutants. Figure 7A shows signal intensities of oncogenic RAS mutants. Figure 7B presents the relative intensities of the different mutants compared to the G12D RAS mutant, which is the most common RAS mutant. By visual inspection, it is easy to note that there are many intersections between these curves that relate the level of predicted RAS signal for each of the oncogenic mutants when expressed at different proportions of total cellular RAS. When two lines cross, the relative ordering of strength of signal produced by the different mutations changes. That is, on one side of the intersection mutant A is stronger than mutant B, but on the other side of the intersection mutant B is stronger than mutant A.

**Figure 7:**
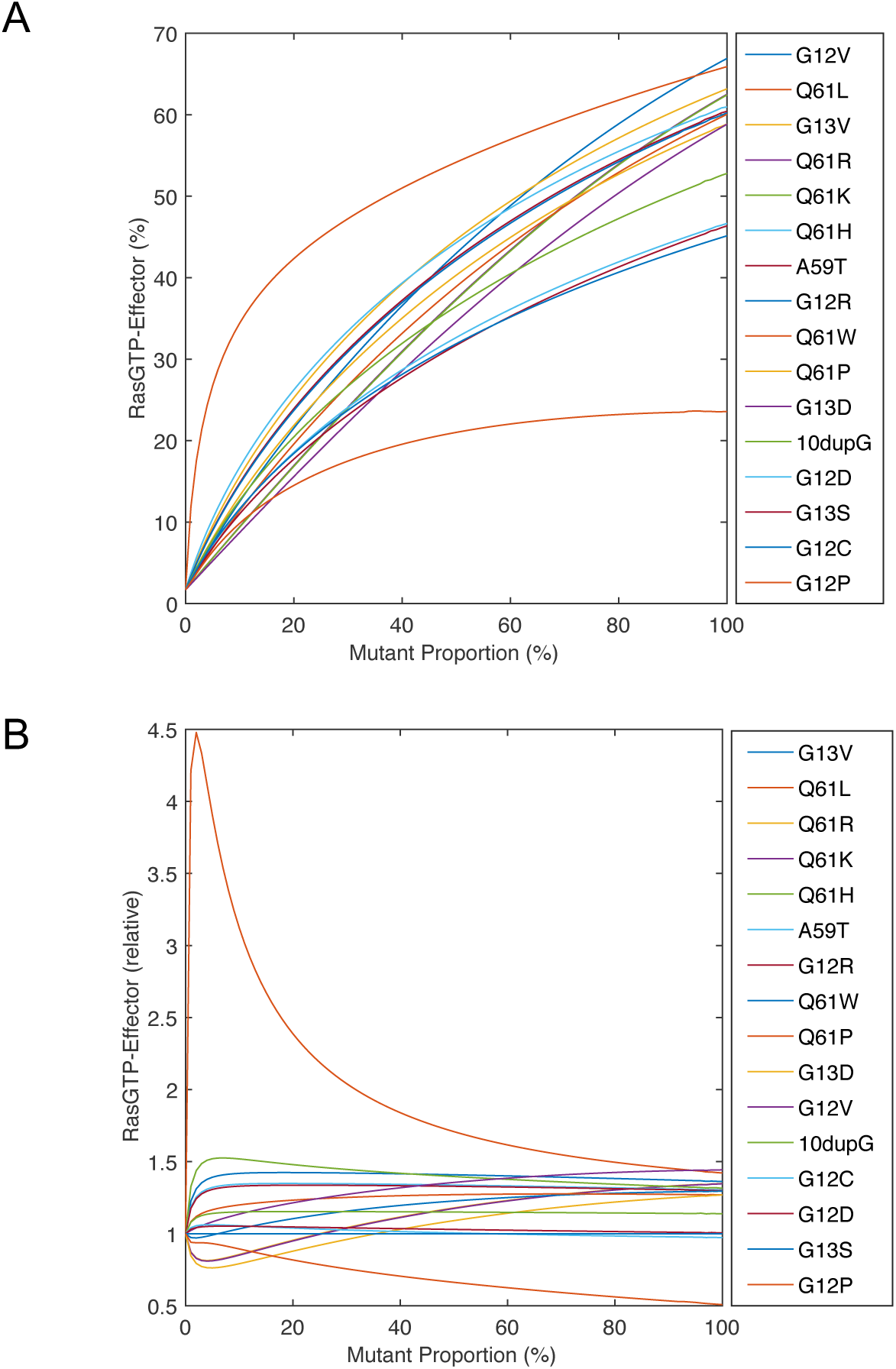
RAS signal as a function of the proportion of total RAS mutated reveals relative strength of oncogenic mutants is not absolute. A) RASGTP-Effector levels for the oncogenic mutants. B) RASGTP-Effector levels for the oncogenic mutants relative to the RASGTP-Effector level for the G12D Ras mutant.

This variation in the relative ranking of oncogenic RAS mutants is highlighted in Figure 8A, which presents the rank order of signal strength as a function of the proportion of total RAS mutated for each of these mutants. These patterns of signaling intensity suggest that the relative strength of two different mutants is not absolute, but is rather context dependent.

**Figure 8:**
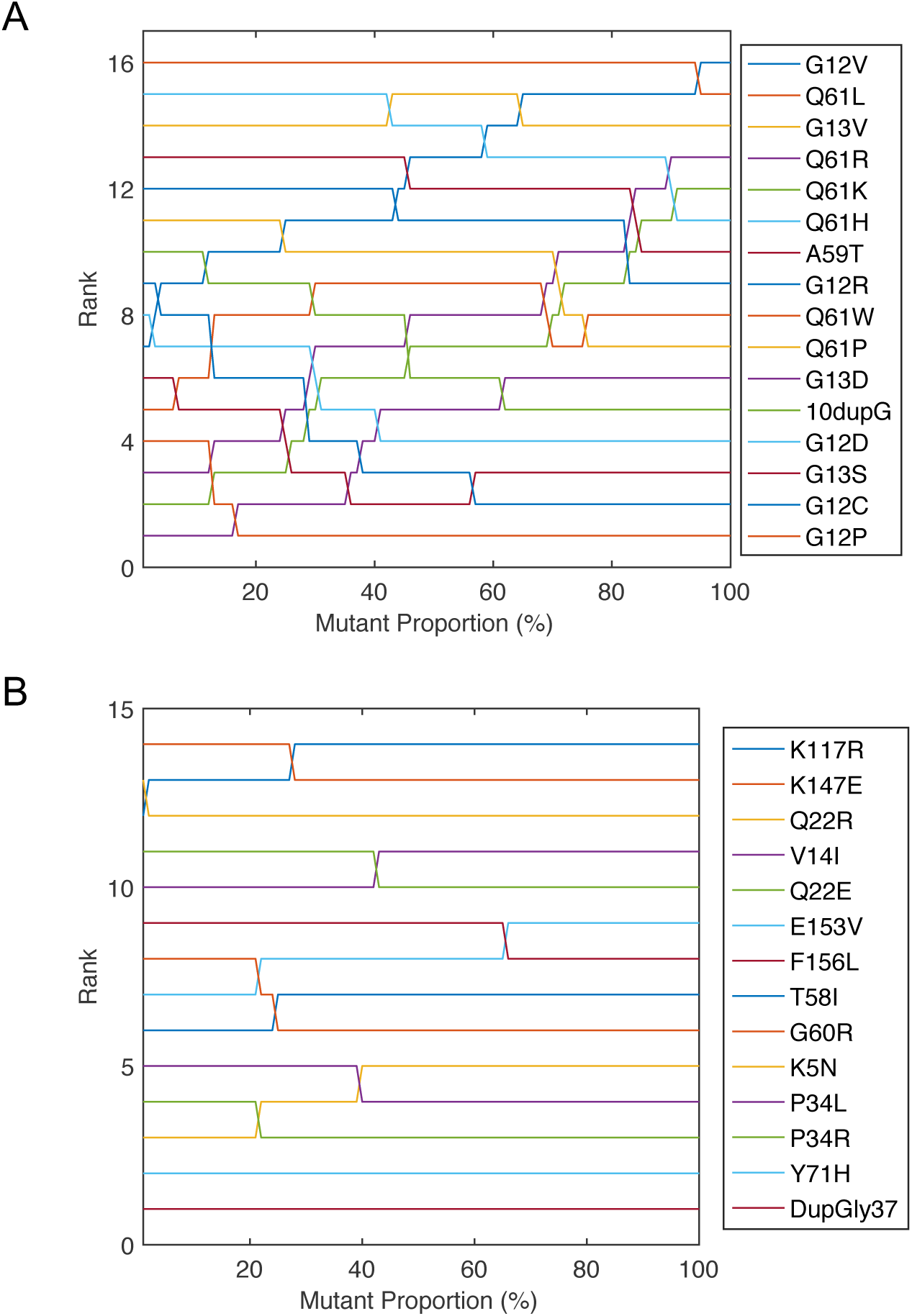
Relative strengths of oncogenic and RASopathy mutants as a function of the proportion of total RAS mutated. A) Oncogenic mutants’ relative strengths. B) RASopathy mutants’ relative strengths.

We also considered the same relationship of signal intensity as a function of proportion of total RAS in the mutant form for the RASopathy mutants. We observe a similar pattern of intersections (Figure 8B). This prediction that the relative strengths of RAS mutants can vary based upon physiological variables such as proportion of total RAS mutated follows naturally from the known biochemical regulation of RAS which is the basis of our mathematical model. Although experimental measurement error and/or gaps in knowledge may shift points of crossing, or even largely shift curves for mutants that were less well characterized, the finding that mutant signal intensity curves will cross seems likely to be a general property of the system.

## DISCUSSION

We have previously used modeling to address specific questions pertaining to RAS mutants G12D, G12V, F28L, G12C, and G13D [8, 15]. Here, an extension of the number of mutants studied was undertaken to develop intuition for how a collection of related, but distinct, pathological mutants might behave. One challenge of modeling pathological mutants is incomplete information. When reaction parameters have not been determined experimentally, we utilized the values of wild-type RAS. (That is, we assume no change.) This assumption is unlikely to be universally true. Still, we argue there are many benefits to such a study. First, consider that our model finds that available data are generally sufficient to predict increased levels of RAS signaling for the majority of these (described) pathological mutants. This suggests that the factors most responsible for pathological activation are generally known.

The analysis also provided new insights. For example, several RASopathy mutants were predicted to have high levels of RASGTP but much lower levels of RASGTP-Effector complex due to their inability to bind effectors well. The prediction that these mutants actually have high RASGTP has, to the best of our knowledge, yet to be tested, but would be an important experiment to investigate RASopathy mutants.

A discrepancy between predictions and observations can suggest that additional biology is at play. Consider that the predicted level of signal for P34R and P34L is quite low, but the level of signal observed has been high [25]. Additionally, consider that the predicted level of signal for D153V and Q22R is moderately increased, but experimentally no change has been noted [25]. Additional biochemical characterization and functional characterization may uncover parameters that result in predictions that match experiments, or perhaps there are additional biological processes that explain the discrepancy. The model has identified interesting candidates for further exploration.

It is also worth noting that mutants Y71H and K5N does not appear to be activated relative to wild-type based on the known biochemical parameters or via experiments [25, 28]. This suggests that there may be yet-to-be-determined mechanisms of activation for these mutants that may be independent of RASGTP or RASGTP-Effector complex. Although low levels of RAS pathway signal were detected experimentally, our computational concurrence strengthens the case that these outlier mutants may be interesting to study more thoroughly.

### A spectrum of RAS strengths

The discovery of germline, intrinsically active, RAS mutations in individuals with RASopathies like Noonan syndrome was initially surprising, because intrinsically active RAS mutations are also associated with cancer. Biochemical characterization of these mutants found them to be less activating than the most common oncogenic RAS mutants. A logical interpretation was that the mutations found in RASopathies were a distinct class that were less activating than the oncogenic mutations, explaining the different disease phenotype.

In years since, the situation has become less clear. Consider that some of the germline mutations in the RASopathy Costello syndrome occur at codon 12 [29] which is also the most common codon mutated in RAS mutations found in cancer [30]. Our analysis demonstrates that the quantity of mutant expressed is an important variable for determining signal strength. Of note, HRAS appears to be expressed at a lower level than KRAS and NRAS in many tissues [26, 27]. Perhaps the critical factor in determining whether a mutation promotes a RASopathy is not the specific mutation, but the quantity of RAS signal generated by the mutation. Strong mutations in less expressed RAS genes may be essentially equivalent to a moderate strength mutation in a more highly expressed RAS gene.

Our computational modeling similarly finds that there is not a clear separation of signal strength between “oncogenic” and “RASopathy” mutants. Clearly, there is a trend that “RASopathy” mutants tend to be weaker, but this difference is not absolute. Further suggesting that there is no clear delineation between oncogenic mutations and RASopathy mutations is the routine detection in cancer genomics of RAS mutations associated with RASopathies [31, 32].Even though these mutations may be comparatively weaker, they still appear capable of promoting cancer.

Overall, this analysis suggests that it may be more correct to think of a wide range of RAS signal intensities that may follow from the nature of the mutant and its expression level, and that downstream phenotypes and disease phenotypes are influenced by the quantity of signal. Recent mouse models investigating dosage of mutation [33, 34] and comparing different mutants [17, 18] can all be interpreted to support the idea that strength of signal, as a function of specific mutation and expression level, is a key factor in determining RAS pathologies.

### Clinical Implications of Variable Rank Order of Strength

Simulations found that the relative strengths of RAS mutants is not fixed, but can vary between different conditions. That is, if one mutant is found to be stronger than another in an experiment, it cannot be assumed that this is always the stronger mutant of the two. It seems reasonable to assume that signal strength is a factor that may contribute to clinically important behaviors of a cancer, such as the response to treatment, rate of tumor growth, and prognosis. Our work here suggests that associations between specific mutations and clinically important behaviors in one cancer cannot be assumed to hold in other cancers.

Variable rank order strength has additional implications for RASopathies. Consider that the RAS mutations in RASopathies are germline and expressed in all tissues, as compared to RAS mutations in cancer which are somatically acquired and only expressed in the tumor and premalignant field. Different tissues within a patient presumably provide different contexts, such as different proportions of total RAS in HRAS, NRAS, and KRAS. The total amount of RAS signal in the different tissues of a RASopathy patient should therefore vary. A different RASopathy patient with a different germline mutations would also be expected to have varying signal strengths between tissues, but the tissues with the strongest signal could vary between patients. This may help explain why different patients with the same disease can have different phenotypes.

This prediction that relative strength varies between contexts could be tested in several different ways. Previously, we have studied RAS signaling by using flow-cytometry to obtain measurements of downstream RAS signaling (measured with anti-pERK antibodies) as a function of RAS mutant expression (using HA-tagged RAS mutant proteins, and anti-HA antibodies to quantify protein expression) [8, 10]. A similar assay, using different RAS mutants, may be able to detect the predicted differences. Additionally, previous experiments expressing different KRAS mutants in the zebrafish pancreas suggested that the capacity of a mutant to activate ERK/MAPK signaling played an important role in cancer development [35]. We hypothesize that if similar experiments were performed where different KRAS mutants were expressed in other tissues, that there would be different patterns of which mutations most promote tumor formation, and that relative pERK induction by the mutants would correlate with these patterns.

## CONCLUSION

There are compelling visions of the future of pathology that involve “computational pathology”, an envisioned sub-discipline of pathology that utilizes computer simulations to help interpret pathological data [36]. However, the visions of computational pathology present few examples and do not address how to extrapolate between pathological mutants. It is difficult to build a field without a foundation. Here, we demonstrate how our computational model of RAS signaling can be used as a tool to study the behavior of pathological mutations and to gain insights into the relationships between mutations and increased signaling (which in turn presumably influences disease phenotypes). We have previously used the model to study problems involving targeted therapies and specific RAS mutations [15], differences in the pathogenicity of different mutants, and methods to target the most common RAS mutants in cancer [8]. We believe that this modeling approach applied more broadly has much to offer computational pathology and personalized medicine.

